# The disease progression and molecular defense response in *Chenopodium quinoa* infected with *Peronospora variabilis*, the causal agent of quinoa downy mildew

**DOI:** 10.1101/607465

**Authors:** Oscar M. Rollano-Peñaloza, Valeria Palma-Encinas, Susanne Widell, Allan G. Rasmusson, Patricia Mollinedo

## Abstract

The downy mildew disease, caused by the biotrophic oomycete *Peronospora variabilis*, is the largest environmental threat to quinoa (*Chenopodium quinoa* Willd.) cultivation in the Andean highlands. However, so far no molecular information on the quinoa-*Peronospora* interaction has been reported. Here, we have developed tools to study the downy mildew disease in quinoa at gene expression level. Living *P. variabilis* could be isolated and maintained in the presence of a fungicide, allowing the characterization of downy mildew disease progression in two differently susceptible quinoa cultivars under controlled conditions. Quinoa gene expression changes induced by *P. variabilis* were analysed by qRT-PCR for quinoa homologues of *Arabidopsis thaliana* pathogen-associated genes. Overall, we observed a slower disease progression and higher tolerance in the quinoa cultivar Kurmi than in the cultivar Maniquena Real. We also observed that quinoa orthologs of *A. thaliana* genes involved in the salicylic acid defense response pathway (*AtCAT2* and *AtEP3*) did not have changes in its gene expression. In contrast, quinoa orthologs of *A. thaliana* gene markers of the induction of the jasmonic acid response pathway (*AtWRKY33* and *AtHSP90*) were significantly induced in plants infected with *P. variabilis*. These genes could be used as defense response markers to select quinoa cultivars that are more tolerant to *P. variabilis* infection.

## Introduction

Quinoa (*Chenopodium quinoa* Willd.) is an allotetraploid annual crop in the amaranth family domesticated by pre-Columbian civilizations in the central Andes of South America approximately 7,000 years ago. Quinoa grains have gained increasing importance in the food market because of its high nutritional value [1, 2]. The ability of quinoa to endure severe drought and high salt concentrations has further raised the interest in quinoa to meet future food demands internationally [1, 3–5]. However, quinoa production in the major original cultivation areas is strongly limited by the downy mildew disease, which can reduce the yield by 35-90% [6–9]. The downy mildew disease is caused by the oomycete *P. variabilis* Gäum. [4, 10, 11] and has been spread to every continent where quinoa is cultivated [12–16]. The worldwide distribution of *P. variabilis* has likely been expanded by commercial trade of infected seeds [17, 18].

*P. variabilis* specifically infects *Chenopodium* species and is an obligate biotroph [19, 20]. Little is known about *P. variabilis* biology, including mode of transmission, leaf penetration and signals for sporangiospore and oospore formation [21]. Most of the studies of *P. variabilis* have been oriented on screening quinoa cultivars for resistance in agricultural fields and observe quantitative differences in susceptibility [6–8, 16, 22, 23]. Some studies have evaluated the resistance of different quinoa cultivars to *P. variabilis* infection under controlled conditions [24, 25] or detached leaf assays [12]. However, mechanistic understanding or molecular studies of the interaction of quinoa with *P. variabilis* or the downy mildew disease progression are not available. With the recent availability of the genomic sequences of quinoa [26–28] and the close relatives to *P. variabilis, Peronospora tabacina* [29] *and Hyaloperonospora arabidopsidis* [30, 31], improved methodology and knowledge on the infection cycle of *P. variabilis* can open up for detailed molecular studies of the quinoa-*Peronospora* pathogenic interaction.

Here, we have developed a method to isolate and maintain *P. variabilis* to facilitate the study of the downy mildew disease in quinoa. We describe the downy mildew disease progression in two quinoa cultivars with different tolerance under controlled conditions. The cultivar with more tolerance upon pathogen attack was selected to identify its defense response mechanisms. The results suggest that quinoa infected with *P. variabilis* expresses defense-related genes that might be involved in the jasmonic acid (JA) pathway (*CqWRKY33* and *CqHSP83*).

## Materials and Methods

### Plant material and growth conditions

Quinoa (*Chenopodium quinoa* Willd.) seeds of the cultivar Maniqueña Real (Real) and Kurmi were kindly supplied by PROINPA (Kiphakiphani, La Paz, Bolivia). Plants were regularly grown and maintained in pots in a greenhouse (Cota Cota, La Paz, Bolivia) under natural light (12 h light / 12 h darkness) and at 17-25°C. Plants were watered three times a week.

### *P. variabilis* isolation and maintenance

Quinoa plants infected with *P. variabilis* were collected from the fields of the PROINPA foundation (Kiphakiphani, Bolivia). Whole infected plants were transplanted *in situ* to pots with fresh soil, covered with plastic bags and transported to our greenhouse. After 24 h, a single-lesion infected leaf was detached from one of the infected quinoa plants and sporangiospores were scraped into sterile ddH_2_O supplemented with 25 µg/ml propiconazole (Propilac 25 EC, Guayaquil, Ecuador). The sporangiospore concentration was adjusted to 1 ×10^6^ per ml and within three hours the suspension was sprayed up to saturation point onto four-week-old quinoa cv. Real plants. Immediately after spraying, semi-transparent polyethylene plastic covers were placed on top of the plants to increase humidity and kept for 24 h. After another 5 days of incubation under greenhouse conditions, plants were covered again for 24 h to favor *P. variabilis* sporulation. For maintenance of the *P. variabilis* isolate, every two weeks, the sporangiospores of a single-lesion infected quinoa leaf were collected into suspension and inoculated onto three-week-old quinoa cv. Real plants as described above, yet without adjusting the sporangiospore concentration.

### Microscopy of *P. variabilis* structures

Staining of hyphae and sporangiospores was performed as described by Koroch, Villani (32), with some modifications. Briefly, quinoa leaves infected with *P. variabilis* were excised in 1 cm^2^ pieces and placed on a microscope slide with the adaxial side facing the slide. Two drops of a solution of I_2_/KI solution (0.5 g I_2_, 1.5 g KI in 25 ml H_2_O) were placed on the abaxial side of the infected leaf, which was incubated at room temperature for 5 min before a cover slip was placed on top. Images were taken with an Optika Vision Pro light microscope (Olympus, Kansas City, Missouri, USA).

### Molecular identification of *P. variabilis*

Total DNA was extracted from *P. variabilis* isolate Kari sporangiospore suspensions using the Purelink genomic DNA Kit according to the instructions of the manufacturer (Thermo Scientific, Carlsbad, CA, USA), with the following modifications: Fresh samples were thoroughly ground under liquid nitrogen in a precooled mortar without letting the samples thaw. Then 600 µl of Purelink Lysis Buffer was added, grinding continued until the samples had thawed, and samples were transferred to 1.5 ml microcentrifuge tubes. DNA was quantified by fluorometry using a Qubit 2.0 Fluorometer (Thermo Scientific).

PCR of the Internal Transcribed Spacer (ITS) region [33] was done with primer pairs DC6/ITS4 [34] and for the *PvCOX2* gene using the primer pairs previously described [35] using the Phusion High-Fidelity PCR Master Mix (Thermo Scientific) supplemented with 0.25 µM of each primer. 20 ng of gDNA was used as template in a 20 µl PCR reaction. The PCR programs had the following settings. One cycle of 98°C for 30 s; 30 cycles of 98°C for 30 s, 50°C for 30 s and 72 °C for 60 s; 1 final cycle of 72°C for 5 min.

PCR products (150 ng) from the ITS region and the *PvCOX2* gene of *P. variabilis* isolate Kari were purified with the QIAquick PCR Purification Kit (Qiagen, Hilden, Germany). The purified PCR products were directly sequenced by the Sanger method (Eurofins, Ebersberg, Germany) and confirmed for the complementary strand.

### Downy mildew disease progression analysis

Three-week-old quinoa plants (cv. Kurmi and Real) were spray-inoculated (to saturation) with either sterile 25 µg/ml propiconazole in ddH_2_O (control) or with a fresh *P. variabilis* sporangiospore suspension [1 ×10^6^ sp/ml] diluted into sterile ddH_2_O and supplemented with propiconazole (to 25 µg/ml). Treated plants were immediately covered with semi-transparent polyethylene plastic bags to raise humidity. The plastic bag covers were left for 24 h. Signs of disease in quinoa leaves were monitored every day. Plants were photographed at 0, 2, 5, 7, 9 and 21 dpi with a digital camera. Leaves were collected at 7 dpi for chlorophyll analysis. The chlorophyll content was estimated from the abaxial side of the second pair of true leaves as described by Liang, Urano (36).

### RNA isolation

Plant tissue from quinoa cv. Kurmi was sampled 48 h post infection (48 hpi) for RNA isolation. One leaf from the second pair of true leaves was cut in half and immediately frozen with liquid nitrogen.

For *P. variabilis* RNA extraction, the sporangiospore/sporangiophore suspensions were prepared by scraping sporangiophores attached to *C. quinoa* leaves from 9 dpi-infected plants. The sporangiospore/sporangiophore suspension was immediately shock-frozen with liquid nitrogen. Total RNA from quinoa or *P. variabilis* was extracted using the Purelink RNA Mini Kit (Thermo Scientific). Briefly, fresh samples were ground under liquid nitrogen in a precooled mortar without letting the samples thaw followed by addition of 1000 µl of Purelink lysis buffer (Thermo Scientific) supplemented with 2-mercaptoethanol [10 µl/ml]. Grinding continued until samples had thawed, and samples were placed in 1.5 ml microcentrifuge tubes. Thereafter, the RNA extraction was performed as described by the Manufacturer.

### Molecular detection of *P. variabilis PvCOX2*

RT-PCR of the *PvCOX2* was done with the primer pairs previously described [35] using the Hot Firepol EvaGreen qPCR Mix Plus (Solis BioDyne, Tartu, Estonia) supplemented with 0.25 µM of each primer. Template was 4 µl of cDNA in a final PCR reaction volume of 20 µl. The PCR program was performed in a LifePro thermocycler (Bioer, Hangzhou, China) and had the following conditions: One cycle of 95°C for 15 min; 30 cycles of 95°C for 30 s, 50°C for 30 s and 72 °C for 60 s); 1 final cycle of 72°C for 5 min. Singularity of PCR products was verified on 2% agarose gels stained with SYBR Safe gel stain (Thermo Scientific).

### cDNA synthesis and qRT-PCR analysis

Isolated RNA was quantified by fluorometry using a Qubit 2.0 and RNA quality was verified by examination of the banding pattern on agarose gels. Synthesis of cDNA was carried out with 500 ng of total RNA added to each 20 µl reaction of the High Capacity cDNA Reverse Transcription Kit (Thermo Scientific). The cDNA samples were stored at −20°C for downstream analysis. qRT-PCR of plant RNA was performed in a StepOnePlus Real-Time PCR system (Thermo Scientific) using Fast SYBR Green Master Mix (Thermo Scientific) supplemented with 0.25 µM of each specific primer and cDNA corresponding to 10 ng of isolated RNA as template. The PCR program had the following conditions: 1 cycle of: 95° C, 20 s; 30 cycles of: (95° C, 15 s; 60°C, 20s; 72 °C, 20s). The specificity of each PCR amplification was determined by melting curve analysis and by analysis in 2% agarose gels. The relative transcript expression was calculated by the Pfaffl algorithm using, *CqACT2* and *CqMON1*, as reference genes. Ten-fold dilutions of cDNA template were used to determine the amplification efficiency for each gene [37].

The primer sequences can be found in Table 1. Primer pairs were designed using Perlprimer [38] so that one of the primers in each pair spanned an exon-exon border, and the primer pairs were checked using Netprimer (premierbiosoft.com) to avoid primer-primer interactions.

**Table 1.**
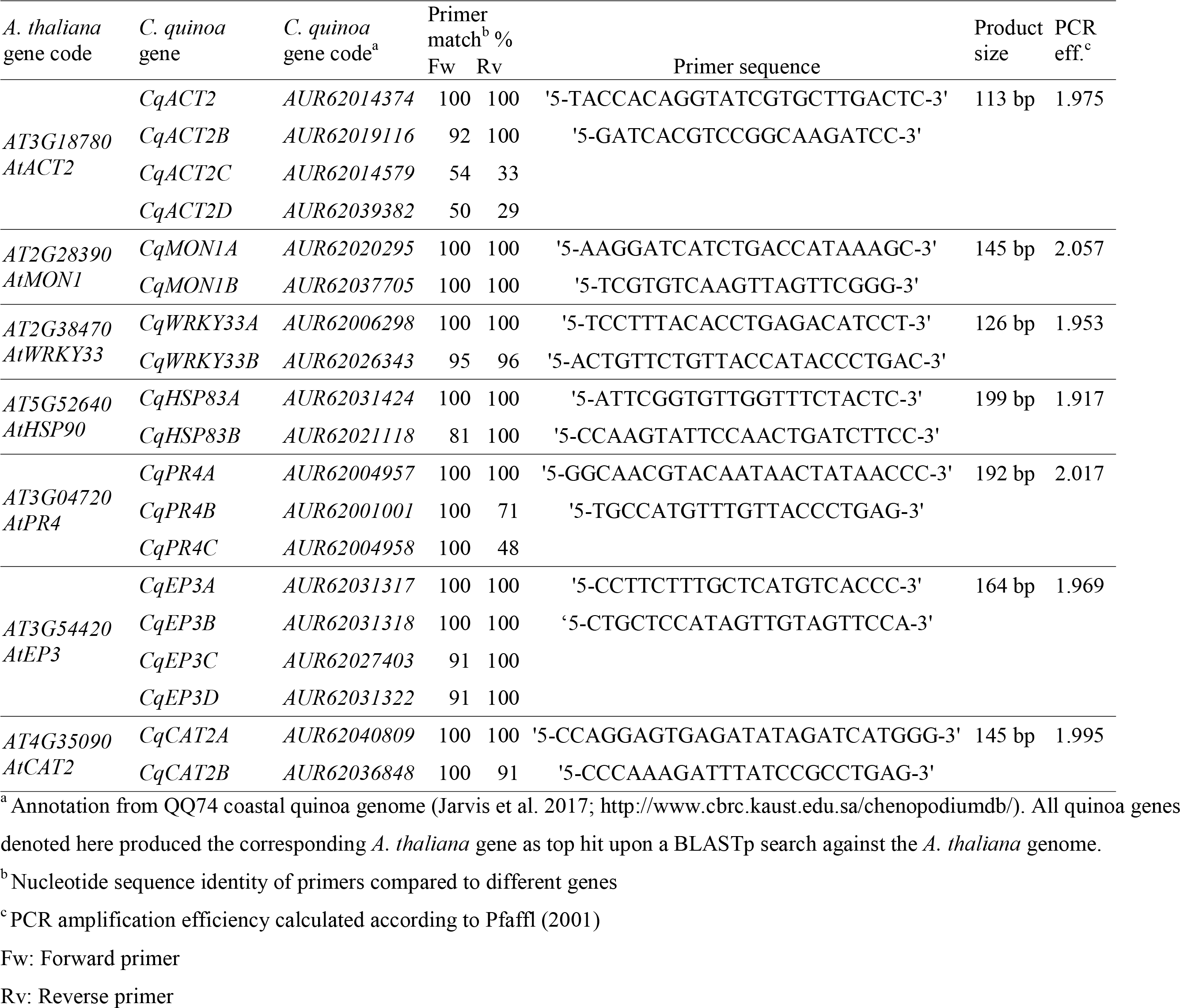
*A. thaliana* genes and orthologs in *C. quinoa* with respective primer sequences for qRT-PCR analysis

### Statistics

Gene expression levels in plants inoculated with *P. variabilis* or mock-treated were compared using Students *t*-test. The statistical analysis was carried out using the R packages plyr [39] and stats [40]. Images were produced using ggplot2 (Wickham 2009).

## Results

### Isolation and characterization of *P. variabilis* isolate Kari

Whole quinoa plants with the abaxial leaves heavily covered with dark grey sporulation structures typical of downy mildew disease were transplanted to pots with fresh soil and transported to our experimental facility greenhouse (Cota cota, La Paz, Bolivia). A single infected leaf was selected and its sporangiospores were scraped off and suspended in sterile ddH_2_O supplemented with the fungicide propiconazole. In preliminary experiments, twenty attempts to produce a virulent sporangiospore suspension without the fungicide were unsuccessful.

The sporangiospore suspension was inoculated onto four-week-old quinoa cv. Real plants within 3 hours of sporangiospore collection. These were covered immediately to increase humidity, and thus increase the likelihood of *P. variabilis* infection and sporulation. Seven days post inoculation (dpi) we observed heavy sporulation in the abaxial part of some leaves. The sporangiospores were collected, suspended and inoculated onto healthy three-week-old quinoa plants. Thereafter, *P. variabilis* was maintained in quinoa cv. Real plants under controlled conditions (13 months at the moment of writing).

In order to verify the identity of *P. variabilis*, we did microscopic observations on the abaxial side of infected quinoa leaves. *P. variabilis* sporulation structures are transparent and difficult to observe on quinoa leaves. However, adaptation of an I_2_/KI staining method allowed observations of typical *P. variabilis* structures at different growth stages (Figs 1A to 1D), including sporangiophores, sporangiospores and oospores. This verified the identity of *P. variabilis*. In order to validate the microscopic observations and preserve the *P. variabilis* isolate identity of this experiment we amplified and sequenced the ITS region and the cytochrome *c* oxidase subunit 2 (*PvCOX2*) gene. The ITS sequence was deposited in the NCBI GenBank under accession number MH999837 and the accession was identified as *P. variabilis*, and we name the isolate Kari. The *PvCOX2* sequence of the isolate Kari was deposited under accession number MK173058.

**Fig 1.**
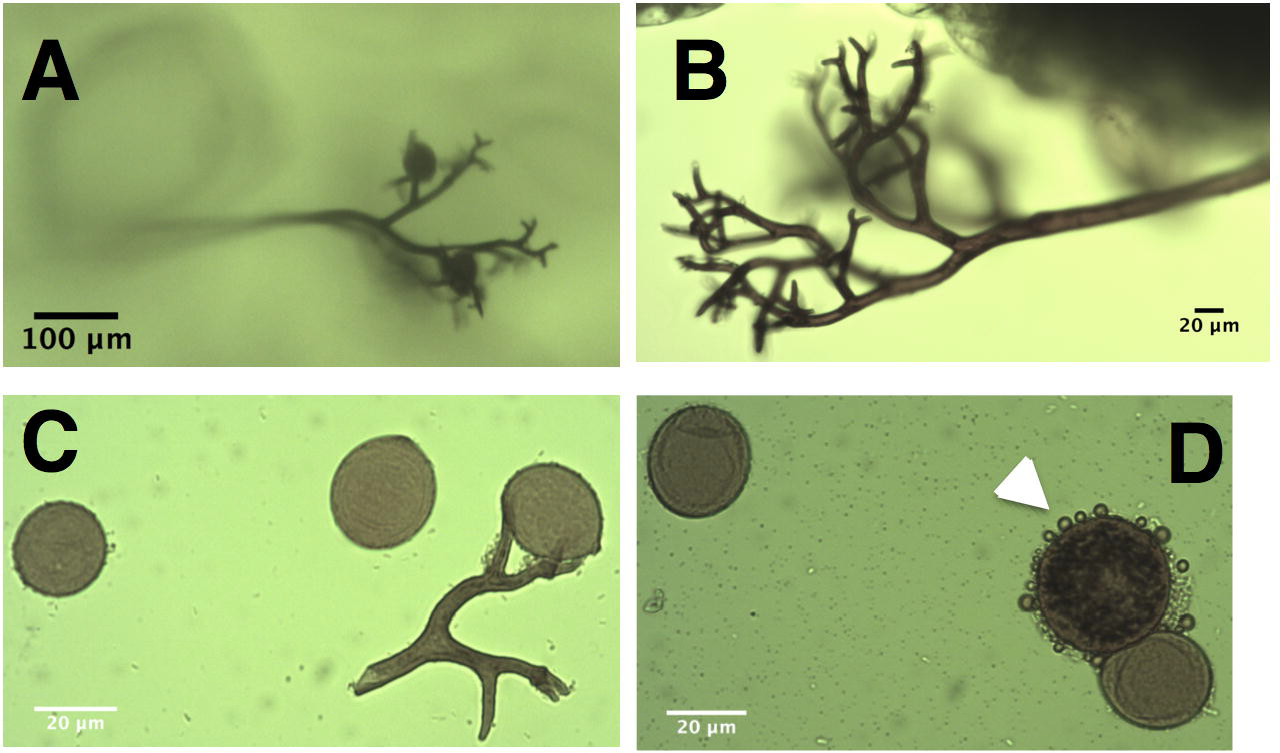
Stained sporangiophores, sporangiospores and oospores of *P. variabilis*. Light microscope images show I_2_’Kl-stained *P. variabilis* structures: (A) sporangiophore branches carrying sporangiospores and growing on the adaxial (top) surface of quinoa leaves; (B) sporangiophores growing out of quinoa leaves; (C) sporangiospores and a broken sporangiophore branch holding sporangia; (D) an oospore (white arrow) next to an sporangiospore.

### Downy mildew disease progression in two *C. quinoa* cultivars

We evaluated the downy mildew disease progression in two quinoa cultivars upon four repeated infection experiments. Three-week-old quinoa plants inoculated with a *P. variabilis* sporangiospore suspension showed folded and moderately chlorotic leaves 5 days after the inoculation (5 dpi), which are initial signs of downy-mildew infection in quinoa [17]. Chlorotic patches of leaves was obvious in the Kurmi cultivar but was barely observed in Real. At 5 dpi none of the cultivars presented sporulation.

Seven days after inoculation (7 dpi) the Real cultivar started to show sporulation signs on the abaxial side of the leaves but Kurmi did not (Fig 2). At this timepoint, chlorotic leaf patches were observed in Kurmi and to a lesser extent in Real (Fig 2). However, image analysis showed that in both quinoa cultivars the chlorophyll content at 7 dpi was significantly lower in leaves of infected plants as compared to control plants (Fig 3).

**Fig 2.**
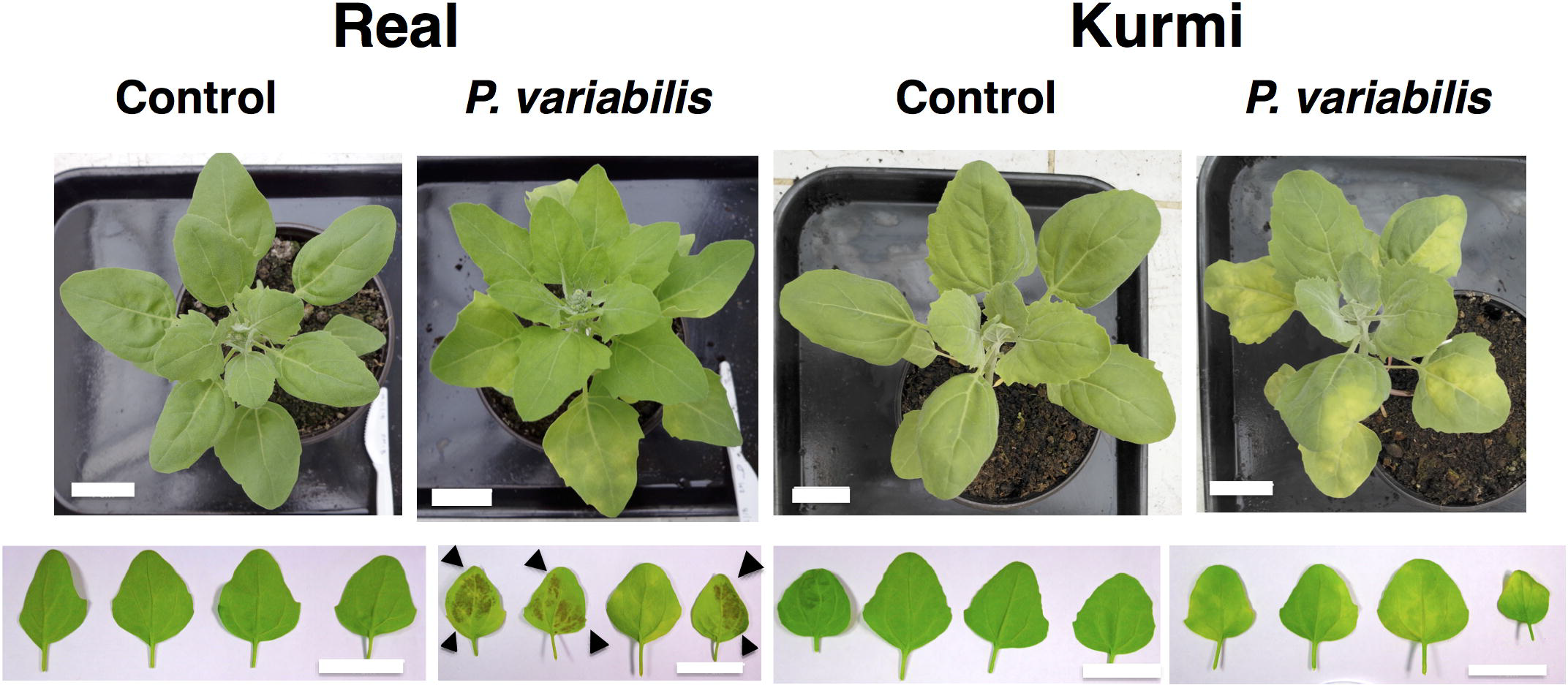
Quinoa plants infected with *P. variabilis*. The figure shows four-week-old quinoa plants 7 days post-infection with *P. variabilis.* Leaf yellowing is shown from the adaxial side (top part) and *P variabilis* sporulation is shown from the abaxial side of leaves from the second pair of true leaves (bottom part). The quinoa cultivar Real displayed sporulation (black arrowheads) but the Kurmi cultivar only chlorosis. The images are representatives of at least three independent experiments, each enclosing three to six biological replicates. All showed similar results. Each leaf in the lower panel was taken from a different biological replicate. The scale bars denote 4 cm.

**Fig 3.**
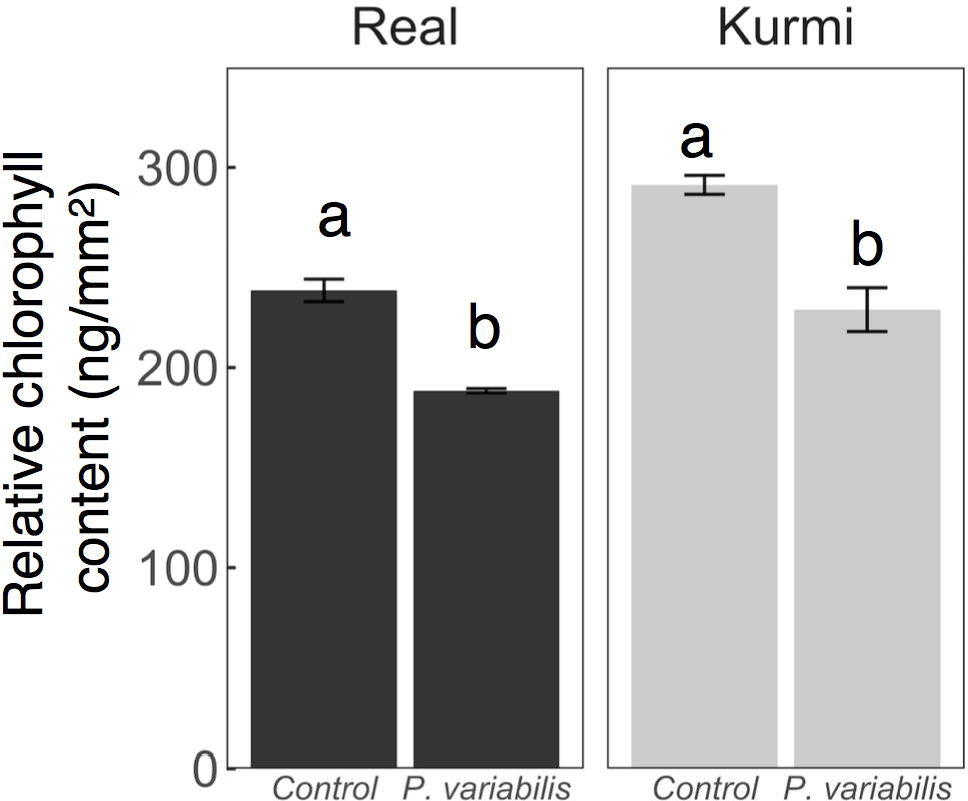
Chlorophyll content in quinoa leaves infected with *P. variabilis*. The second true leaf of three-week-old quinoa plants at 7 dpi was collected and analysed. Data shows means ± SE per treatment (n = 4).

The effect on vitality of quinoa was further evaluated until 21 dpi (Fig 4). Here, we observed that from 9 dpi, both Kurmi and Real showed heavy sporulation on the abaxial part of the leaves. In the absence of pathogen, the Real cultivar grew faster than Kurmi. Real plants infected with *P. variabilis* were severely and negatively affected as compared to the mock-treated plants (Figs 4A and 4C). Real infected with *P. variabilis* showed sporulation in most of its leaves and many leaves were wilting or fully necrotic at 21 dpi (Fig 4C). In contrast, Kurmi displayed infected leaves, but the sporulation was localized only to the chlorotic parts of the leaves (Fig 4D). In both Kurmi and Real, new leaves emerged without signs of infection. However, in Real the leaves were derived mostly from side-branches emerging by a loss in apical dominance, whereas in Kurmi they emerged from both the main stem and from side-branches (Figs 4C and 4D). The Real cultivar infected with *P. variabilis* also induced early flowering (Fig 4C) as compared to its mock-treated counterpart (Fig 4A). Early flowering was not observed in the Kurmi cultivar (Figs 4B and 4D). No necrotic lesions with defined edges were observed at any time point. Overall, the results suggested that the Kurmi cultivar has an ability to delay sporulation, withstand the infection, and form new leaves outgrowing the infection, and thus being more tolerant to *P. variabilis* infection than Real.

**Fig 4.**
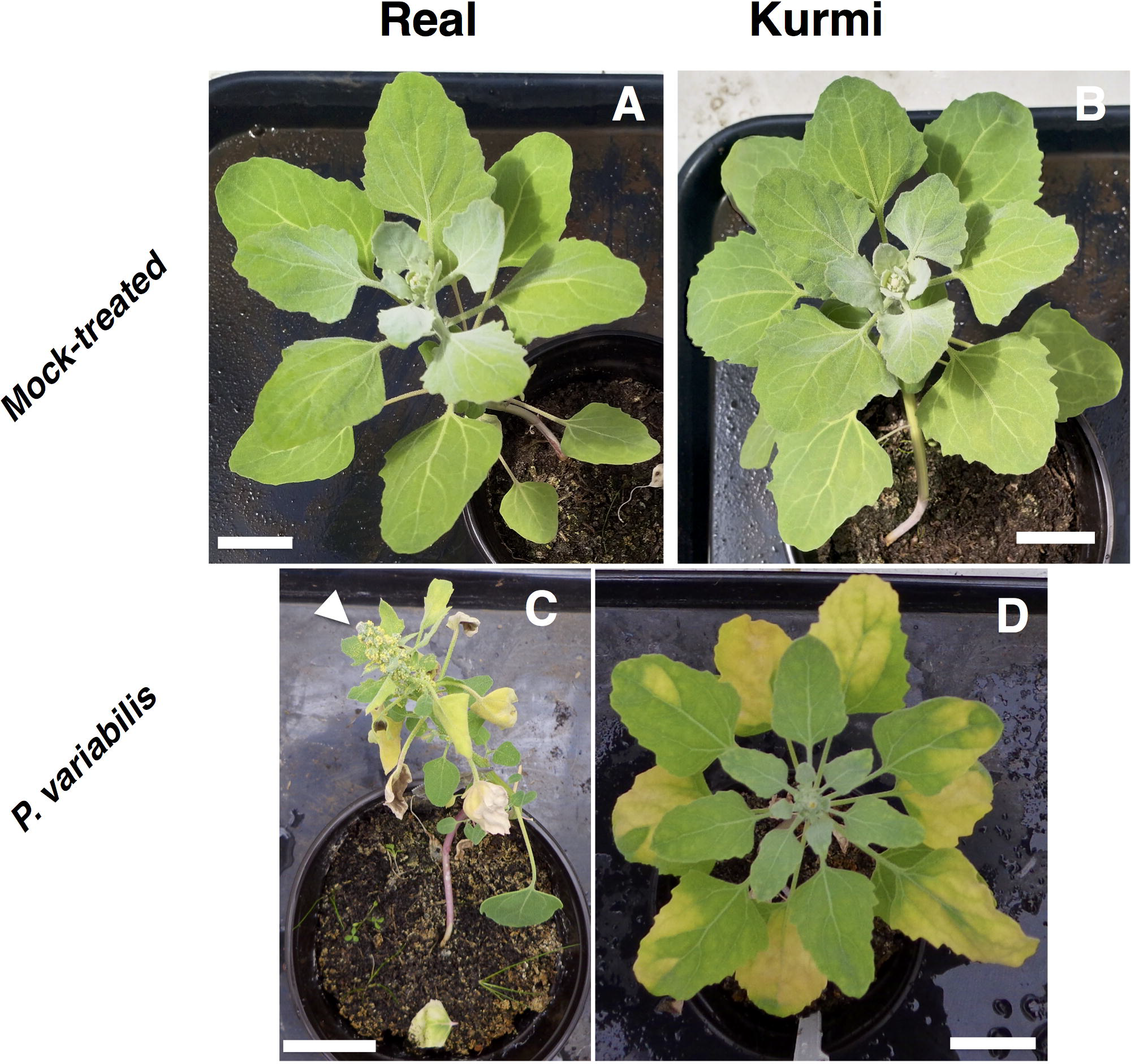
Six-week-old quinoa plants 21 days after infection with *P. variabilis*. The figure shows the symptoms of the downy mildew disease (C and D) as compared to mock-treated controls (A and B) in quinoa cv. Real (A and C) and cv. Kurmi (B and D). The images are representatives of two independent experiments of each three biological replicates, and with similar results. The white arrowhead points at the early flowering observed in the infected Real cultivar.

### Quinoa gene expression defense response against *P. variabilis*

To investigate the increased tolerance of Kurmi we performed gene expression analysis of the defense response against the pathogen. Quinoa plants did not shown any infection signs during the first 2 days after inoculation. In order to verify that plants had become infected by the treatment at the time of sampling (2 dpi) for gene expression analysis, we performed molecular detection of *P. variabilis* in RNA samples. RT-PCR with primers against the *P. variabilis* cytochrome *c* oxidase subunit 2 gene (*PvCOX2*) produced a 600 bp product in plants infected with *P. variabilis* at 2dpi, but no PCR product was observed in mock-treated plants (Fig 5). This verified that quinoa plants were infected with *P. variabilis* as intended.

**Fig 5.**
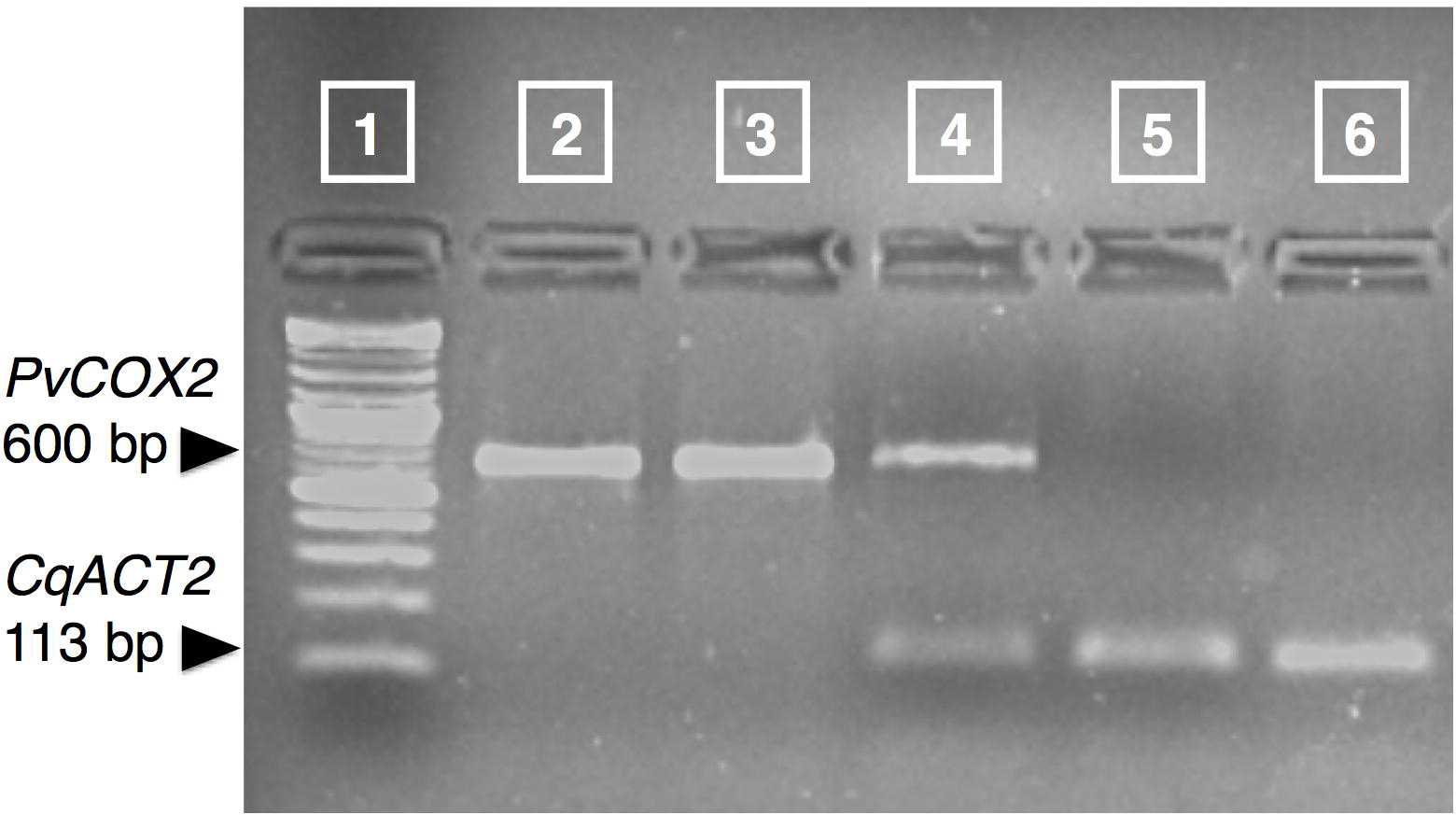
Detection of *P. variabilis* by RT-PCR against the cytochrome *c* oxidase subunit 2 gene. Lane 1: 2-log DNA ladder (New England Biolabs); lane 2: *P. variabilis* genomic DNA; lane 3: *P. variabilis* cDNA; lane 4: cDNA derived from a plant infected with *P. variabilis;* lane 5: cDNA derived from a mock-treated plant; lane 6: C. *quinoa* genomic DNA. As a reference, the amplification of the plant control gene *(CqACT2)* is shown.

Based on *A. thaliana* microarray data for expressional stability under stress [41], putative quinoa reference genes for mRNA quantification were selected. We verified the presence of *A. thaliana* gene orthologs in quinoa by two-way BLASTp [42] searches against the quinoa genome. For most of the genes in *A. thaliana* there are at least two corresponding homologs in quinoa, because quinoa is allotetraploid [26]. Therefore, the top hit from each forward BLAST search was selected for primer design. The gene selected was identified with a letter at the end of the gene name abbreviation in order of sequence similarity to the *A. thaliana* gene (Table 1). The reference genes selected were the orthologs of *At3g18780* (*Actin 2, AtACT2)* and *At2g28390 (Monensin Sensitivity 1, AtMON1)*. The quinoa ortholog of *AtACT2* with the highest BLAST score out of 4 ortholog sequences was *AUR62014374* (identified in our study as *CqACT2*). *AtMON1* had two quinoa orthologs: *AUR62020295* and *AUR62037705* (identified as *CqMON1A* and *CqMON1B*, respectively). Due to the high nucleotide sequence identity (98%) between the *CqMON1* orthologs our primer pair targeted both genes, and we denote both genes as *CqMON1* (Table 1). The selected quinoa reference genes showed similar stability in qRT-PCR product formation over the population of samples, displaying average Ct and standard deviation of 20 ± 0.6 and 23 ± 0.5 for *CqACT2* and *CqMON1, respectively*. *CqACT2* was selected as the primary reference gene due to its higher expression levels. The results were verified by *CqMON1*, which showed similar expression with and without *P. variabilis* treatment (Fig. 6A).

**Fig 6.**
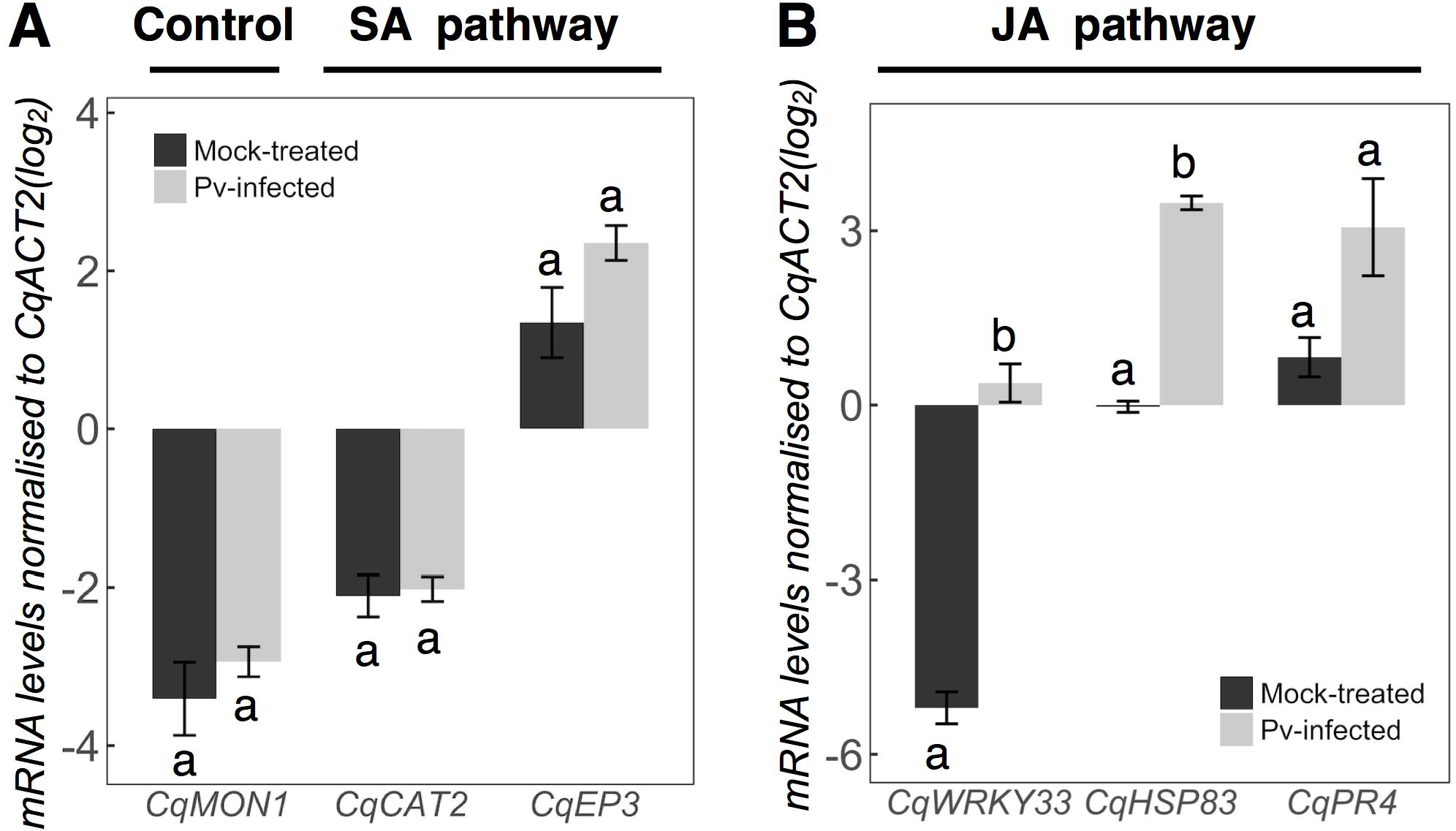
Gene expression in quinoa plants in response to *P. variabilis*. The figure shows defense response genes orthologs of quinoa cv. Kurmi at 48 h after spray-inoculation with *P. variabilis* or mock control. (A) The control gene (*CqMONJ*) and orthologs of genes involved in the SA pathway; (B) Orthologs of genes involved in the JA pathway. Significant difference (p<0.05) between mock-treated and infected plants is denoted by different letters.

Plant defense response against biotrophs usually involves the activation of the salicylic acid (SA) signaling cascade in order to activate the hypersensitive response and halt the biotrophic pathogen growth [43]. Therefore, we tested the quinoa orthologs of the *AtCAT2* [44] and *AtEP3* [45] genes, which are both active in the *A. thaliana* SA response. For more studied SA pathway genes, as *AtNPR1* (*AT1G64280*), clear orthologs could not be identified in quinoa. Both *CqCAT2A* and *CqEP3* (Table 1) were found to accumulate mRNA to similar levels in plants infected with *P. variabilis* and in mock-infected plants at 48 hpi (Fig. 6A).

The SA pathway can be antagonized by the JA signaling cascade upon recognition of necrotrophic pathogens [46]. The JA pathway triggers the synthesis of plant defense proteins and phytoalexins but also often down-regulate the hypersensitive response [47]. Plants can also trigger the JA pathway defense response upon recognition of biotrophic pathogens. Therefore, we assessed the mRNA abundance of quinoa orthologs of *A. thaliana* genes induced by the JA defense response pathway: *AtHSP90* [48], *AtWRKY33* [49] and *AtPR4* [50]. At 48 hpi, all three quinoa orthologs (*CqHSP83*, *CqWRKY33* and *CqPR4)* displayed elevated mRNA abundance values in plants infected with *P. variabilis*, yet only the signals for *CqHSP83 (p = 2×10^−5^)* and *CqWRKY33 (p = 2×10^−4^*) were significantly different from that in the mock-treated plants (Fig. 6B).

## Discussion

The novel *P. variabilis* isolate Kari had reproductive structures and produced disease symptoms (Fig. 1) similar to other isolates described before [17, 21]. The fungicide propiconazole was a key factor for the successful isolation of infectious *P. variabilis*, indicating that the fungicide inhibits the growth of fungi that otherwise can parasitize on oomycetes like *P. variabilis*. Specific growth inhibition in fungi but not in oomycetes can be achieved because propiconazole inhibits one of the steps in the synthesis of ergosterol, the major sterol in fungi [51]. In contrast, *Peronospora* and other Peronosporales (Oomycetes) do not synthesize ergosterol and thus contain no target for the fungicide [52]. Sequencing of the ITS region verified that our strain isolated from the Bolivian Andean plateau belongs to the *P. variabilis* species [11]. The ITS region in *P. variabilis* is not enough to determine the geographic origin of isolated strains [17], yet the sequence of *CqCOX2* confirmed its origin in South America.

*P. variabilis* isolate Kari was compatible with both quinoa cultivars, Real and Kurmi (Fig. 2 and 4). The downy mildew disease symptoms produced by *P. variabilis* isolate Kari were chlorosis, foliar curling, and heavy sporulation on the abaxial side of the leaves. These symptoms are consistent with the downy mildew disease symptoms in susceptible quinoa cultivars observed in the agricultural fields [9, 17].

Growth and development of the Real cultivar was more affected by *P. variabilis* than the Kurmi cultivar (Fig. 4). The early flowering in quinoa cv. Real produced by *P. variabilis* is a typical symptom of stress-induced flowering [53]. Stress-induced flowering has for example been observed in *A. thaliana* infected with the oomycete *H. arabidopsidis* [54]. The overall better growth and higher developmental similarity between control and infected plants observed in Kurmi (Fig. 2 and 4), indicate that Kurmi is more tolerant to *P. variabilis* infection than Real. This is consistent with the high resistance to downy mildew observed in the Kurmi cultivar compared to Real in cultivations of quinoa in the Andean plateau (A. Bonifacio; *personal communication*). Therefore, we suggest Kurmi is a suitable candidate to study quinoa defense response mechanisms at molecular level.

The quinoa cultivar Kurmi did not show signs that would suggest that hypersensitive response (HR) was triggered after infection (Fig. 2 and 4). HR is normally triggered by the SA defense response pathway when plants interact with biotrophic pathogens [43]. We found that quinoa orthologs of the *A. thaliana* genes involved in the SA defense response pathway were neither induced nor repressed (Fig. 6A). For example, *AtCAT2* that encodes a putative functional catalase [55] is suppressed by SA to increase hydrogen peroxide levels and eventually trigger HR [44]. In our results we did not observe significant changes in its ortholog gene expression (*CqCAT2*, Fig. 6A). Similarly, the chitinase *CaEP3* of the Chenopodiaceae *Chenopodium amaranticolor* has been observed to be expressed under biotic stress or by SA [45], but the ortholog in quinoa *CqEP3* was not significantly changed in our experiments (Fig. 6A). The results thus suggest that the SA pathway defense response would have not been induced in the quinoa cultivars studied, consistent with the lack of hypersensitive response.

Instead, chlorosis signs were observed in infected leaves where the pathogen was visibly sporulating from the abaxial side of the leaf in both quinoa cultivars. The chlorosis was visibly stronger and more spatially variegated in the Kurmi cultivar than in Real (Fig. 2 and 4). Similar results were observed in *A. thaliana* susceptible varieties (compatible interactions) in response to the infection with the biotrophic pathogen *H. arabidopsidis* [56]. Chlorosis can be stimulated by methyl-jasmonate and can be a signal that the JA-mediated defense response pathway is activated [57–59]. Some of the known molecular markers of the JA-mediated defense response in *A. thaliana* and grapevine (*Vitis vinifera*) are the transcription factors *AtWRKY33* and *VvWRKY33*, respectively [49, 60, 61]. In our results the quinoa *CqWRKY33* gene was significantly induced and this could mean that the JA-mediated defense response was induced (Fig. 6B). Although the JA defense response pathway has been usually known to be the main defense response against necrotrophic pathogens [43], recent studies revealed that JA may also play a role in the defense response against biotrophic pathogens [31, 62]. The activation of JA-mediated defense response in quinoa upon infection with *P. variabilis*, can be supported by the induction of the quinoa pathogenesis-related protein 4 (*CqPR4*) and *CqHSP83* (Fig. 6B). Orthologs of *CqPR4* as the *A. thaliana AtPR4* [50], the *PR4* in sugarcane (known as *Sugarwin*) [63] and the *ZmPR4* in maize [64] are known to be induced by treatment with Methyl Jasmonate (Me-JA). Consistently, the *A. thaliana* heat shock protein 90 (*AtHSP90*) [65–67] together with its ortholog in *Nicotiana benthamiana* (*NbHSP90*) [68] are the closest orthologs of *CqHSP83* and they are known to have a central role in the JA defense response pathway.

Quinoa cv. Kurmi gene expression in response to the oomycete *P. variabilis* displayed similarities to the *A. thaliana* Col-0 response to *H. arabidopsidis* Waco9 (compatible interaction). *AtHSP90*, *AtWRKY33* and *AtPR4* involved in the JA pathway were differentially expressed in *A. thaliana* plants after 3 days of infection with *H. arabidopsidis* but the genes involved in the SA pathway *AtEP3* and *AtCAT2* were not [31]. Overall, the results suggest that at least parts of the JA defense response pathway, but not the SA pathway, is activated in quinoa upon recognition of *P. variabilis.* Still, despite the response, and the higher tolerance to *P. variabilis* infection in Kurmi than in the Real cultivar, Kurmi should be designated a semi-susceptible cultivar.

It is important to notice that the degree of *P. variabilis* compatibility reported here for Kurmi and Real might change with a different isolate of *P. variabilis*. This compatibility between plants and pathogens can be cultivar- and isolate-specific, as it has been described in the plant model *A. thaliana* interacting with *H. arabidopsidis* [69]. Given the documented high genetic diversity of quinoa in the Andean highlands [70], we may also expect a high genetic diversity of *P. variabilis*, possibly with different compatibilities with the different quinoa cultivars.

In conclusion, both quinoa cultivars were susceptible to infection by the novel *P. variabilis* isolate Kari. The infection has stronger and more rapid effects over the vitality of Real, showing higher proportion of dead leaves, reduced growth and altered morphology as compared to Kurmi. Further, none of the cultivars presented signs suggesting that hypersensitive response had been triggered, consistent with the molecular data that suggested that the SA defense response pathway was not activated. The differentially induced genes *CqWRKY33* and *CqHSP83*, orthologs of genes known to be involved in the JA defense response pathway in other species, suggest that the defense response of the quinoa cultivar Kurmi against *P. variabilis* isolate Kari might be mediated by the JA signaling pathway.

Understanding the molecular response and defense mechanisms involved in the higher *P. variabilis* tolerance during infection in the Kurmi cultivar can contribute to the development of resistant quinoa cultivars in future breeding programs.

## Acknowledgements

We are grateful to Proinpa Institute (Kiphakiphani, Bolivia) for the generous donation of quinoa seeds and quinoa plants infected with *P. variabilis*.

